# Stretched-Exponential Modeling of Anomalous T_1*ρ*_ and T_2_ Relaxation in the Intervertebral Disc *In Vivo*

**DOI:** 10.1101/2020.05.21.109785

**Authors:** Robert L. Wilson, Leah Bowen, Woong Kim, David A. Reiter, Corey P. Neu

## Abstract

**Purpose:** Intervertebral disc degeneration (IVDD), resulting in the depletion of hydrophilic glycosaminoglycans (GAGs) located in the nucleus pulposus (NP), can lead to debilitating neck and back pain. Magnetic Resonance Imaging (MRI) is a promising means of IVD assessment due to the sensitivity of MRI tissue relaxation properties to matrix composition. Furthermore, anomalous (i.e. non-monoexponential) relaxation models have shown higher sensitivity to specific matrix components compared to conventional monoexponential models. Here, we extend the use of the stretched exponential model, an anomalous relaxation model, to IVD relaxometry data.

**Theory and Methods:** T_1*ρ*_ and T_2_ relaxation data were measured in the cervical IVDs of healthy volunteers and IVDs adjacent to cervical fusion, and analyzed using both conventional and stretched-exponential (SE) models. Model differences were evaluated via goodness of fit in the healthy data. Normalized histograms of the resultant quantitative MRI (qMRI) maps were described using stable distributions, and data were compared across adjacent disc segments.

**Results:** In the healthy IVDs, we found lower mean squared error in the SE relaxation model fitting behavior compared to monoexponential models, supporting anomalous relaxation behavior in healthy IVDs. SE model parameter α_T1*ρ*_ increased level-wise in the caudal direction, especially in the nucleus pulposus, while conventional T_1*ρ*_ and T_2_ monoexponential measures did not vary. For IVDs adjacent to cervical fusion, SE parameters deviated near the fusion site compared with those in the healthy population.

**Conclusion:** SE modeling of T_1*ρ*_ relaxation provides greater sensitivity to level-wise variation in IVD matrix properties compared with conventional relaxation modeling, and could provide improved sensitivity to early stages of IVD degeneration. The improved model fit and correlation between the SE α_T1*ρ*_ parameter with IVD level suggests SE modeling may be a more sensitive method for detection of GAG content variation.

## INTRODUCTION

Intervertebral disc degeneration (IVDD) is a common condition, affecting over 80 million adults in the United states alone, which can lead to debilitating neck and back pain (1,2). The IVD structure consists of the collagen-heavy ring-shaped annulus fibrosus (AF) and the glycosaminoglycan (GAG)-rich nucleus pulposus (NP). The IVD is vertically bound first via thin layers of hyaline cartilage and then by the vertebrae themselves (3). IVD resistance to external load is dependent on the hydrophilic GAG molecules that maintain IVD hydrostatic pressure. In IVDs positioned in increasingly caudal locations, the NP increases in volume to withstand greater loads by producing increased hydrostatic pressure within the IVD. While the exact etiology of disc degeneration remains unknown, depletion of GAGs in the NP is associated with IVD degeneration (4).

Magnetic Resonance Imaging (MRI) is a promising tool for IVD assessment. GAG deficiencies are routinely assessed in the research setting using histology (5–8). However, it has long been a goal of medicine to detect early soft tissue degeneration *in vivo* (9,10). MRI is an ideal modality for early detection of IVDD *in vivo* due to its excellent noninvasive soft tissue contrast and routine clinical use. Furthermore, MRI’s ability to provide enhanced tissue characterization through quantitative relaxometry sets it apart as a diagnostic tool. In a comparable tissue, T_2_ relaxation analysis has been heavily utilized for *in vivo* cartilage assessment due to its sensitivity to collagen and water content (11–13). In contrast, while T_2_ values in the IVD have primarily been shown to be sensitive to water content, T_1*ρ*_ values have been shown to be sensitive to GAG content (14–19).

While conventional relaxometry measures are helpful tools to characterize IVD structure, monoexponential T_1*ρ*_ and T_2_ values have shown limited specificity to individual matrix components (20). In contrast, anomalous relaxation has been observed in similar biological systems with non-monoexponential signal models showing additional tissue composition information (21,22). The stretched-exponential (SE) function has been used widely for modeling biological and physical phenomenon (23,24). Recently, a SE function, which may more appropriately reflect relaxation time distributions associated with different tissue compartments and compositions, was used to model T_2_ relaxation data of bovine nasal cartilage (21). The resulting improved decay fit was attributed to a strong correlation between the stretching parameter *α* and GAG content *in situ,* suggesting an improved specificity to GAG content over conventional relaxation methods.

Our objective was to extend and apply a stretched-exponential relaxometry model to characterize IVD structure *in vivo.* Conventional and SE models of T_1*ρ*_ and T_2_ relaxation data were compared in the adjacent cervical IVDs of healthy patients, and additionally in a patient that had recently undergone a cervical fusion procedure. We demonstrate a wider dynamic range in SE T_1*ρ*_ model parameters across disc level, suggesting greater sensitivity to known level-wise compositional variations between healthy IVDs compared to conventional *in vivo* measures. Additionally, SE analysis from IVDs adjacent to cervical fusion suggest changes in IVD composition consistent with degeneration. Improved sensitivity gains achieved through the SE model could lead to improved biomarkers, allowing for detection of subtle tissue composition changes associated with IVDD in adjacent discs following cervical fusion.

## THEORY

The SE model for fitting transverse magnetization (T_2_) decay profiles has previously been derived and discussed in detail (21,25). The conventional monoexponential T_2_ relaxation decay equation can be written:

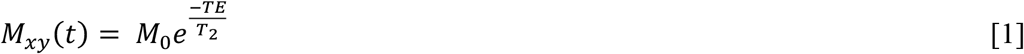

Where *M_xy_* is the transverse magnetization, *M*_0_ is the initial magnetization, T_2_ is the calculated time constant, and *TE* is the echo time. The stretched-exponential decay equation can be written:

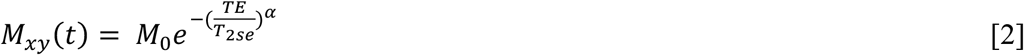

where the stretching parameter, α, allows for the modeling of broad continuous distributions of relaxation times suitable for capturing varying degrees of tissue microstructural complexity.

T_2_ relaxation times in complex tissues can depend on the selection of interpulse delays due to a variety of effects (e.g. diffusion through field inhomogeneities and spin-spin coupling). The influence of these effects, which result in varying degrees of spin dephasing, can be minimized by using a sufficiently short interpulse delay. In the limit of vanishing interpulse delay time relative to pulse length, magnetization is effectively locked in the rotating frame, and T_2_ relaxation approaches the spin-lattice relaxation time in the rotating frame, T_1*ρ*_ (26). Through varying the application of the spin-lock pulse frequency, relaxation in the rotating frame can be measured over a wide range of frequencies below the Larmor frequency, permitting the observation of interactions between water and extracellular matrix molecules (i.e. exchange of protons between mobile matrix proteins and water) (16). As T_2_ and T_1*ρ*_ probe similar motional properties, and T_2_ has shown anomalous decay in IVDs (3), we hypothesized that T_1*ρ*_ may also exhibit anomalous decay. The conventional model for monoexponential T_1*ρ*_ relaxation can be written:

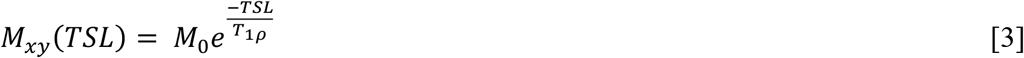

where *TSL* is the duration of the spin lock pulse. Similar to Equation 2, the stretched-exponential T_1*ρ*_ relaxation model can be written:

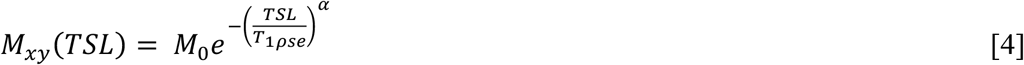

Using equations 1–4, two different models (conventional and SE) for spin-spin and spin-lattice relaxation in the rotating frame (T_2_ and T_1*ρ*_) yield six different model parameters to analyze: T_2Mono_, T_2SE_, α_T2_, T_1*ρ*Mono_, T_1*ρ*SE_, and α_T1*ρ*_.

## METHODS

Institutional Review Board approval was obtained from the Human Research Protection Program at Purdue University and Emory University for all aspects of this study. The methods described were carried out in accordance with the approved guidelines, and informed consent was obtained from all participants.

### Raw Data Acquisition and Processing - Imaging in Adjacent IVDs of Healthy Subjects

Cervical IVDs (C2C3-C6C7) of 15 healthy subjects (7/8 males/females; average age = 24.7) were imaged using a 3T GE MRI scanner. A magnetization-prepared, angle-modulated and partitioned *k*-space spoiled gradient echo snapshots (MAPSS) T_1*ρ*_ sequence was acquired with the following parameters: spin lock power: 500Hz, spin lock times: [1, 5, 20, 40, 60] ms (27). A T_2_ sequence was also acquired (echo times: [6.78,13.97,21.15,42.72] ms). Shared imaging sequence parameters were: FOV: 14×14 cm^2^, Matrix: 256×128 px^2^, Slice Thickness: 4 mm, Views Per Segment: 64, TR 1.2 s, Number of Slices: 26, ARC Acceleration Factor: 2, NA: 4.

For T_1*ρ*_ relaxation, the first spin lock time (TSL) data was excluded from relaxation fitting due to uncharacteristically low signal at the first time point resulting in a substantial increase in mean squared error (MSE) for all models compared with removal of the data. Voxelwise processing was chosen for analysis because of its relatively low noise compared to other techniques (Supplemental Figure 1). Decay curve fitting for both the conventional and SE models were completed utilizing a standard curve fitting toolbox (MATLAB) with bounds on model parameters (T_2_, T_1ρ_: [0,120] | α bounds: [0.4,1]). Signal-to-noise ratios (SNRs) were calculated by comparing the average value of the whole disc signal to the mean of a background sample equal in pixels from the final TSL/TE of each disc. Discs with low SNR (<2) and individual pixels with relaxation decay fits with an *R*^2^<0.66 (28) or with an outlier decay time (Grubb’s Test, α=0.05) were excluded from the study. Overfitting was evaluated via residual scatter plots and histogram analysis (Supplemental Figure 2). Quantitative MRI (qMRI) maps of the outputs from conventional (i.e. T_2Mono_ and T_1*ρ*Mono_) and SE fits (i.e. T_2SE_, α_T2_, T_1*ρ*SE_, and α_T1*ρ*_) were created based on voxel-wise fitting (Figure 2B). The qMRI maps were smoothed with a locally weighted scatterplot smoothing (LOWESS) filter (span: 10 pixels) for noise and edge effect minimization (29).

**Figure 1:**
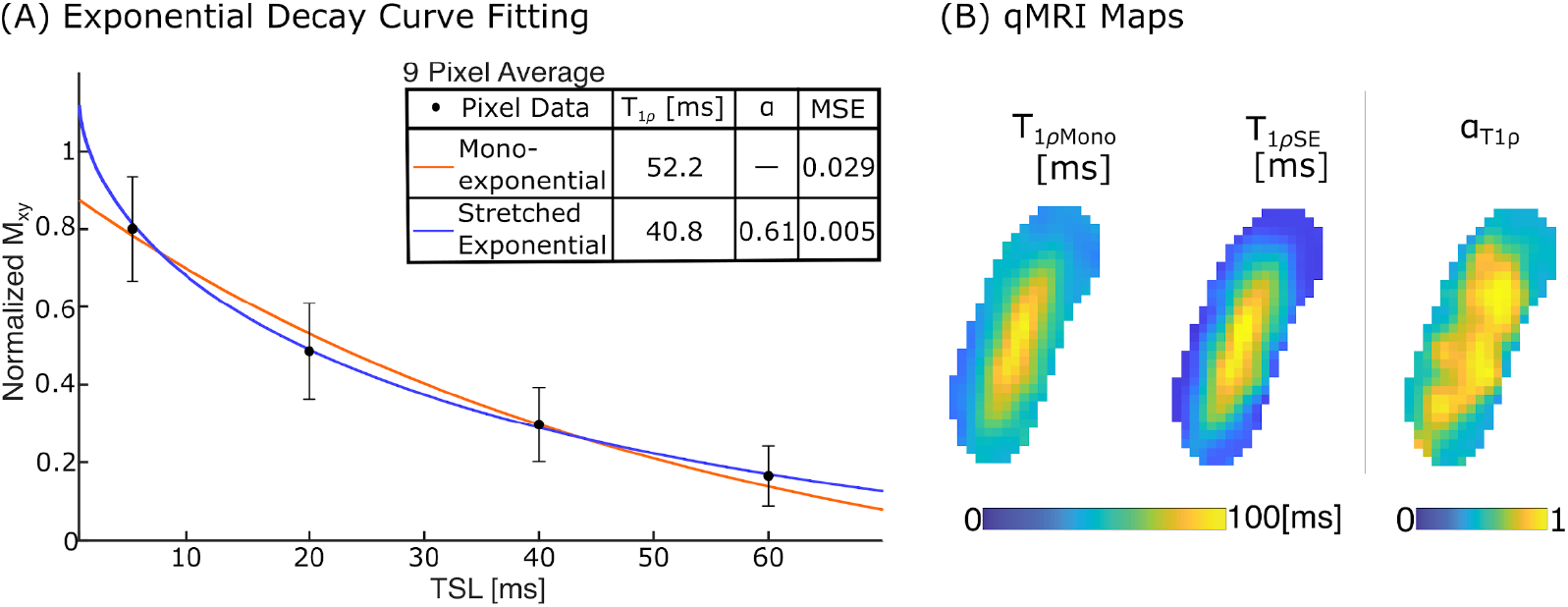
The exponential decay curve model fit reveals an increase in spatial heterogeneity for the stretched-exponential T_1*ρ*_ and α_T1*ρ*_ compared to conventional T_1*ρ*_. (A) The decay curves (a 9-pixel average is shown here for clarity) are qualitatively similar, yet the resulting T_1*ρ*_ values for both models are drastically different. The inclusion of the α value results in a significantly smaller MSE (*p*<0.001). (B) Higher levels of detail are visible in the T_1*ρ*SE_ and α_T1*ρ*_ maps compared to the monoexponential map. This sensitivity may lead to increased sensitivity to matrix composition differences.

**Figure 2:**
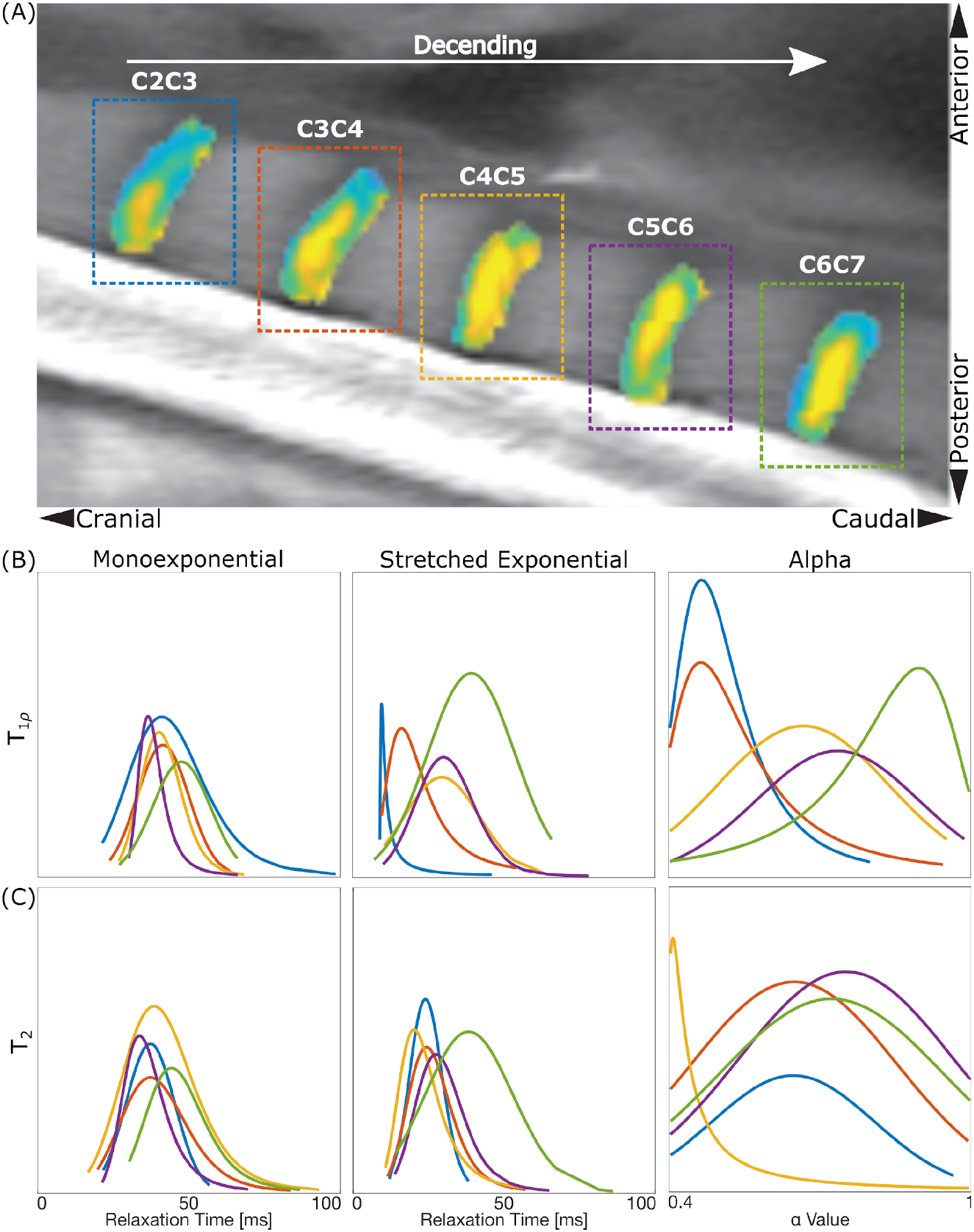
In a single subject, T_1*ρ*_ and T_2_ SE distribution of parameter fit values within each disc level. T_1*ρ*_ SE data suggest a monotonic shift in the distribution peak position with disc level. (A) The qMRI maps, overlaid on an anatomical image (C2C3-C6C7), show the positional relationships of the discs. (B,C) The single subject T_1*ρ*_ and T_2_ stable distributions show a cranial-caudal relationship in the stretched-exponential and alpha terms, while the monoexponential terms do not.

### Raw Data Acquisition and Processing – Imaging of Adjacent IVDs following Cervical Fusion

Images were acquired in the sagittal plane of the cervical spine from a single subject (Female, Age: 46) five months following anterior cervical discectomy and fusion surgery at C6-C7 using a 3T Siemens PrismaFit scanner. T_1*ρ*_ weighted images were acquired using a magnetization prepared spoiled gradient echo prototype sequence (30) with imaging parameters: spin lock power: 400 Hz, spin lock times: [0, 10, 20, 30, 40, 50, 60] ms, FOV: 22×22 cm^2^, Matrix: 256×256 px^2^, Slice Thickness: 3 mm, TR: 9.1 s, Number of Slices: 16, NA: 4. Decay curve fitting was achieved via the same process as the healthy subject data (i.e. voxelwise via MATLAB’s curve fitting toolbox).

### Analysis of Regions of Interest in Multiple Adjacent IVDs

Regions of interest (ROIs) were manually segmented using the T_1*ρ*_ weighted image with the shortest TSL, showing good soft tissue contrast, to create binary masks containing the whole disc, the AF and the NP (in healthy subjects only). To characterize differences in relaxation values from a given ROI, qMRI map histograms were created and normalized by the total number of pixels and fit with stable distributions with standard bounds (α = (0,2], β = [-1,1], γ = (0,∞), δ = (-∞,∞)) as the distributions were found to be non-normal (*p*>0.05). Analogous to the position of a normal distribution being represented by the distribution mean, the stable distribution position is represented by the δ value with α and β representing the distribution symmetry and skewness, respectively.

For model parameter comparison between disc levels, herein termed a *level-wise analysis,* differences in the resultant δ values of each model parameter’s stable distribution (i.e., for each distribution representing histograms of monoexponential, SE, and α data) were analyzed as a function of anatomical disc level (Supplemental Table 1). For visual comparison (see Figure 3C, Figure 4C, and Figure 5C-D), each subject’s IVD δ values were normalized by subtracting the lower fitting bound (LB) and then dividing by the bound fitting range (upper fitting bound (UB) - LB) and offsetting all IVDs per subject by the C2C3 δ value (δ_C2C3_):

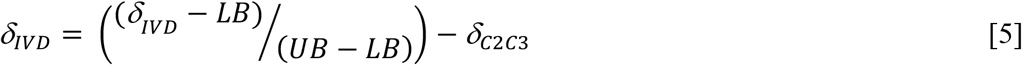

**Figure 3:**
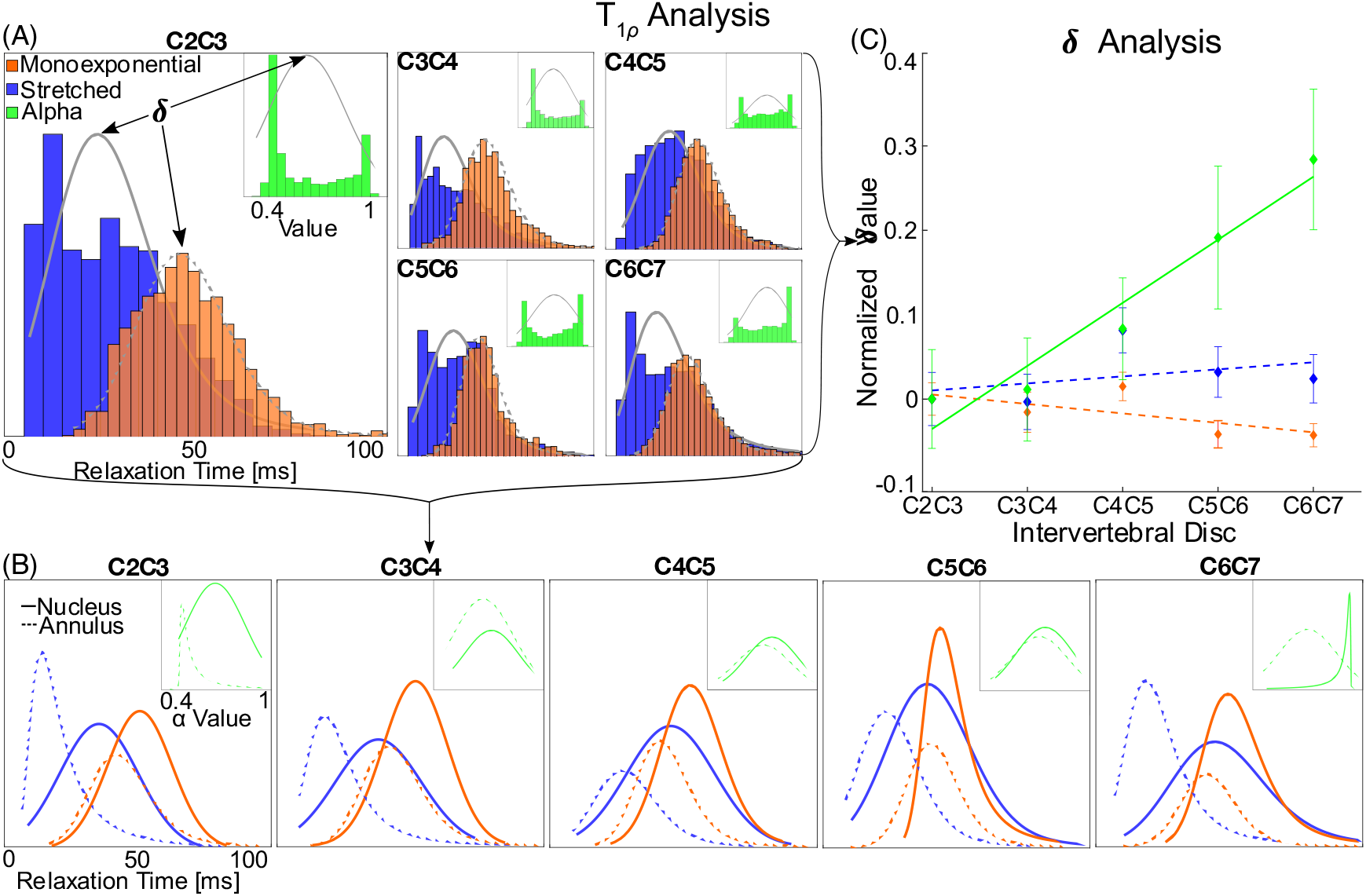
The T_1*ρ*_ stable distribution analysis shows a significant monotonic increasing relationship between IVD level and peak values for the SE α term, but not for either time constant. (A) The population-level (*n*=15) T_1*ρ*_ stable distributions caudally increase. Arrows indicate the peak (δ) value of each distribution. (B) Population-level analysis of the nucleus pulposus (NP) versus annulus fibrosus (AF) indicates a greater change in peak values for the NP compared to the AF, particularly in the α data. (C) The level-wise Spearman’s rank correlation is significant for the α term (*p*=0.0029) but not significant for either T_1*ρ*_ metric (Monoexponential: *p*=0.120, SE: *p*=0.32). Solid regression lines indicate *p*<0.01; Dashed lines represents no significance. Error bars represent standard error of the mean (SEM).

**Figure 4:**
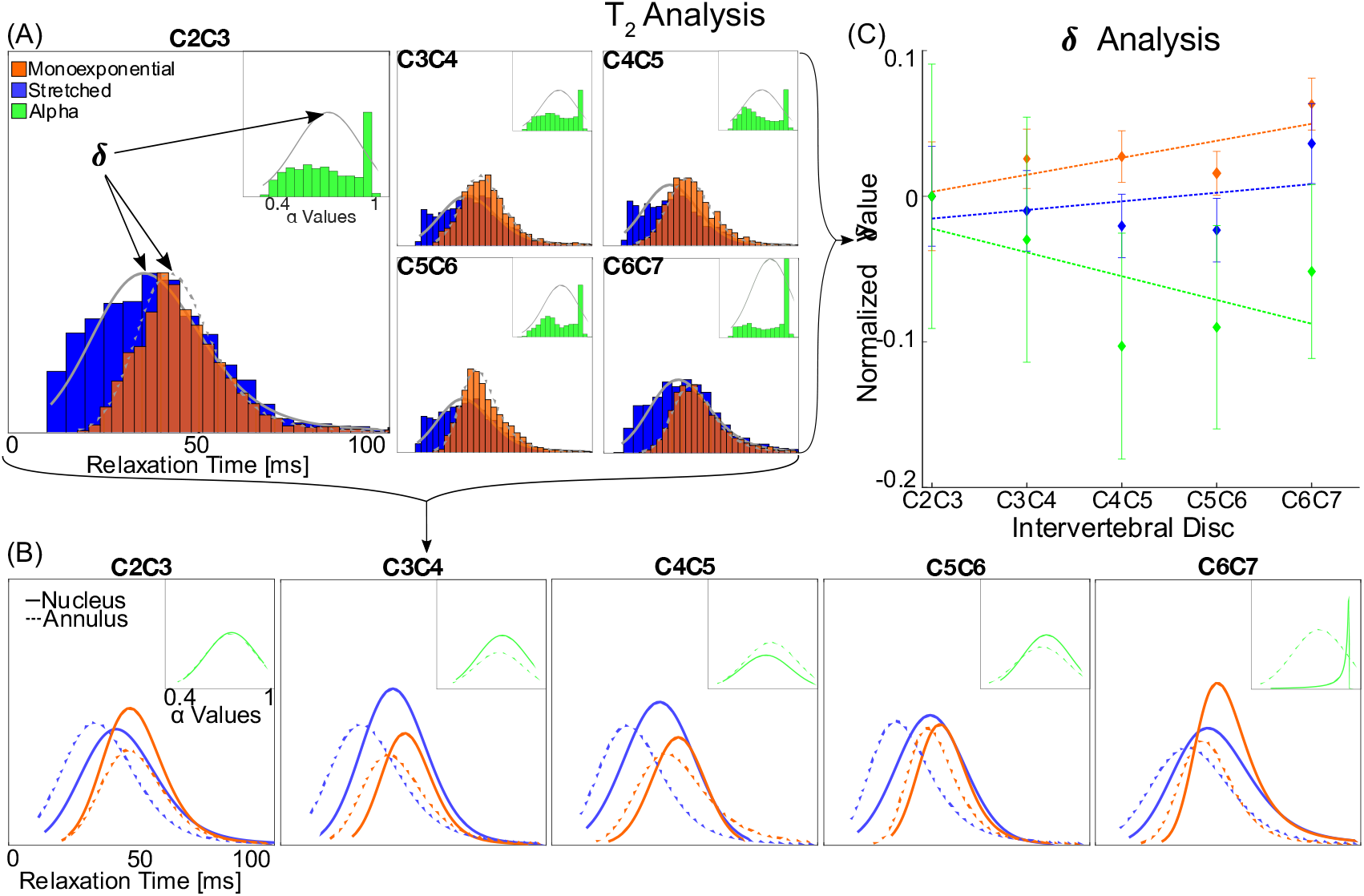
The T_2_ stable distribution analysis shows no correlation between distribution peak values (δ) and IVD level. (A) The population-level (*n*=15) T_2_ stable distributions have no position-dependent correlation. Arrows indicate the peak value of each distribution. (B) Population-level analysis of the nucleus pulposus (NP) versus annulus fibrosus (AF) indicates no detectable change in peak values for the NP compared to the AF. (C) The level-wise Spearman’s rank correlations found no significance (Monoexponential: *p*=0.19, SE: *p*=0.43, α: *p*=0.31). Dashed regression lines represent no significance. Error bars represent SEM.

### Statistics

Model fit comparisons were made based on the MSE of both models using a one-way ANOVA. Histogram normality was assessed using Shapiro-Wilks test. Absolute interdisc differences were determined using Kruskal-Wallis. The level-wise comparison of IVD levels was evaluated using a Spearman’s rank correlation coefficient testing for monotonic relationships (Supplemental Table 1). Statistical significance was set at *p*<0.05.

## RESULTS

In the healthy IVD, we evaluated goodness of fit between models and found lower mean squared error in the SE compared to monoexponential T_1*ρ*_ relaxation model (Figure 1, Supplemental Figure 1). Single subject level-wise analysis showed an increase in SE T_1*ρ*_ model parameters moving caudally with IVD segment, but monoexponential models revealed no qualitative differences (Figure 2). MSEs derived from model fits were lower in the stretched-exponential model than the conventional model for both T_1*ρ*_ and T_2_ (*p*<0.001) with similar levels of fit (Supplemental Figure 2). Level-wise variation was more prominent in the T_1*ρ*_ parameters than T_2_. While the T_1*ρ*Mono_ and T_1*ρ*SE_ level-wise analyses qualitatively indicate a relationship, a stronger correlation existed in the α_T1*ρ*_ analysis. Level-wise comparison of α (α_T1*ρ*_, α_T2_) suggests a monotonic variation caudally (C5C6-C6C7), particularly in the NP compared to the AF (Supplemental Figure 3).

The population level (*n*=15) T_1*ρ*_ stable distribution analysis demonstrated a significant relationship between IVD level and the SE model α_T1*ρ*_ parameter (Figure 3A, Supplemental Table 1). Caudally, the T_1*ρ*Mono_ and T_1*ρ*SE_ data remained relatively constant in the level-wise analysis, while the α_T1*ρ*_ indicated level-wise increase. α_T1*ρ*_ values in the NP increased dramatically in the caudal direction whereas the annulus fibrosis data showed a more moderate level-wise dependence (Figure 3B, Supplemental Figure 4). The T_1*ρ*Mono_ and T_1*ρ*SE_ data showed no discernable trend for each disc component. The whole disc normalized level-wise analysis (Eqn. 5) revealed a statistically significant monotonic relationship between IVD level and α_T1*ρ*_ (Figure 3C) (Spearman’s Correlations: *p*=0.003 | ρ=0.404) while the TipMono and T_1*ρ*SE_ δ values did not (T_1*ρ*Mono_: *p*=0.197 | T_1*ρ*SE_: *p*=0.322).

The population level analysis of T_2_ data showed no significant correlation between parameters (T_2Mono_, T_2SE_, and α_T2_) and IVD segment. All model parameters remained relatively constant by level (Figure 4A). No major differences were found when analyzing NP and AF separately (Figure 4B). No correlation was found between disc level and T_2_ values (Figure 4C) (T_2Mono_: *p*=0.190, T_2SE_: *p*=0.425, α_T2_: *p*=0.312).

T_1*ρ*_ parameter maps were generated in a patient post-fusion and level-wise statistics were calculated for levels (C2C3-C4C5) adjacent to implanted hardware (Figure 5A-B). The monoexponential T_1*ρ*_ level-wise analysis was consistent with the trends calculated from healthy subjects data (Figure 5C). T_1*ρ*SE_ and α_T1*ρ*_ analysis showed similar values to healthy subjects for C2C3-C3C4, but sharply deviated from the healthy trends at C4C5 (Figure 5C-D), i.e. in IVD segments most adjacent to implant hardware.

**Figure 5:**
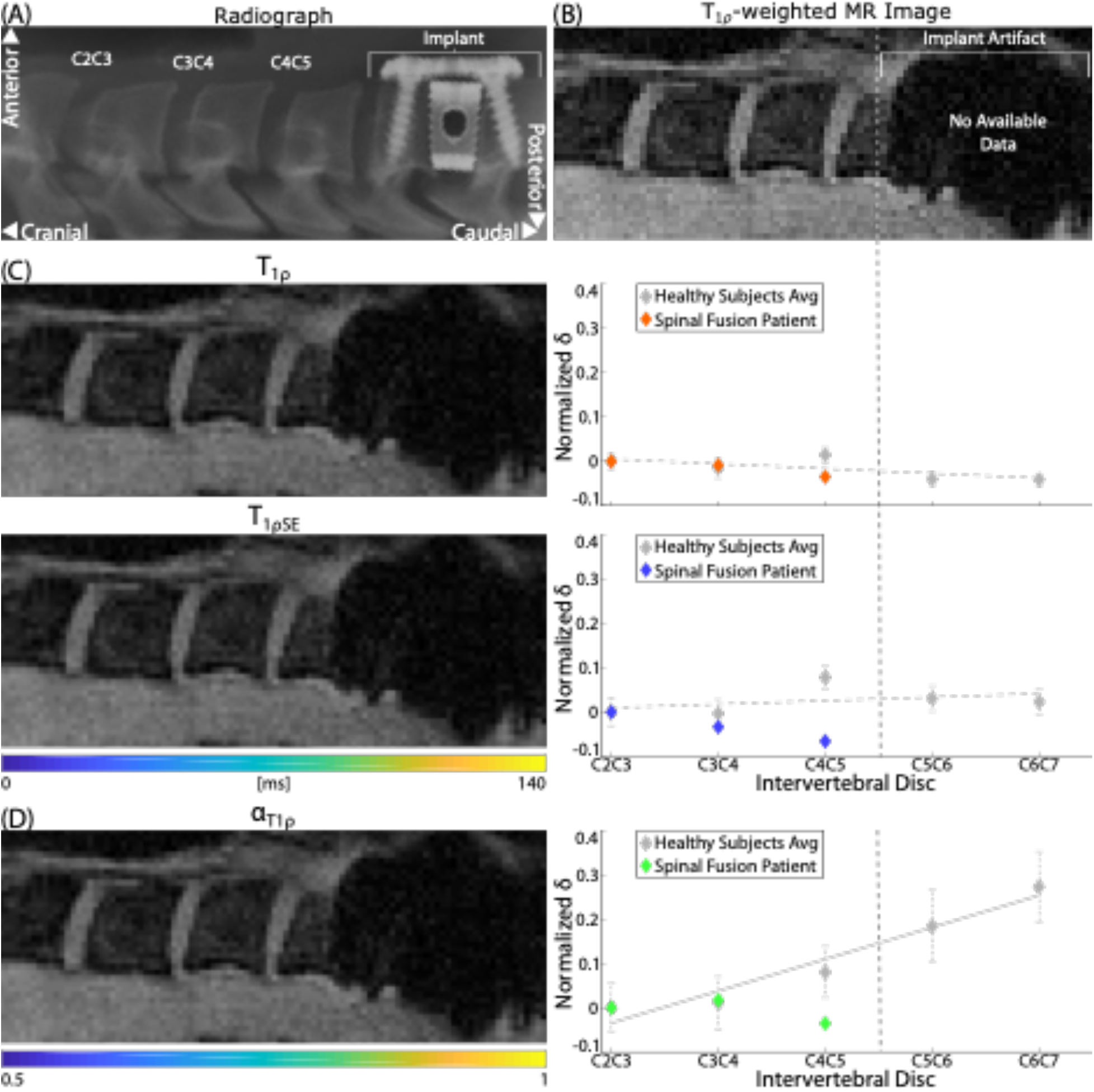
In IVD segments adjacent to cervical fusion, the stretched-exponential model parameters deviate from the calculated trends observed in healthy subjects, indicating compromised IVD structure near the implant site. (A) Radiograph of the post-operation implant hardware for cervical fusion, and adjacent IVDs. (B) A T_1*ρ*_-weighted MR image of the post-fused patient demonstrates the imaging artifact created by the implant, which does not preclude analysis of adjacent IVD segments. (C) The monoexponential (orange) and stretched-exponential (blue) T_1*ρ*_ values from the post-fusion patient overlaid on the healthy subject data (grey; repeated also from Figure 4C). The T_1*ρ*_ data indicates no level-wise differences while T_1*ρ*SE_ shows a deviation near the site of fusion compared with that in healthy individuals. (D) Comparison of α_T1*ρ*_ (green) with healthy individuals shows a deviation in parameter values near the level of fusion, suggestive of structural differences in C4C5.

## DISCUSSION

The purpose of this study was to explore the use of the SE model to investigate cervical IVD properties in healthy subjects and following fusion *in vivo.* T_1*ρ*_ and T_2_ relaxation models were investigated. Overall, we found α_T1*ρ*_ significantly correlated with disc level in the healthy population. Both monoexponential models and the SE T_2_ model did not show any disc level dependence in these subjects. SE analysis of T_1*ρ*_ images following cervical IVD fusion, suggests a higher sensitivity to degenerative changes in IVD matrix integrity compared with traditional relaxation modeling. These preliminary results support the possibility of effectively evaluating changes during IVDD using anomalous T_1*ρ*_ modeling.

One interpretation of the stretched-exponential model is its sensitivity to the presence of microscopic heterogeneity manifesting in decay time distributions. In the molecular tumbling autocorrelation framework, monoexponential relaxation follows naturally from spins in a homogeneous environment like that observed in solutions NMR (31). When fitting both models to bulk water, the time constants for the monoexponential and SE models are expected to be equivalent (Figure 6) with an α value of one in the SE model indicating a lack of sample complexity that would give rise to multiple exponential behavior. However, with increasing tissue complexity, the interaction of water spins with their environment (i.e. GAG and other proteins) modulated by the tissue microenvironment (e.g. in structure, pH, chemical exchange), causes non-monoexponential decay rates and thus shift toward anomalous behavior. Moreover, this anomalous behavior is expected to further change with alterations in IVD structure in the pathogenesis of degeneration, e.g. in IVDs adjacent to cervical fusion (Figure 5). Additionally, the slower molecular interaction of larger molecules in complex tissues, particularly GAGs and the associated water content, may play a primary role in anomalous relaxation behavior, especially reflected in the α parameter, which demonstrates the largest dependence on IVD level (Figure 3).

**Figure 6:**
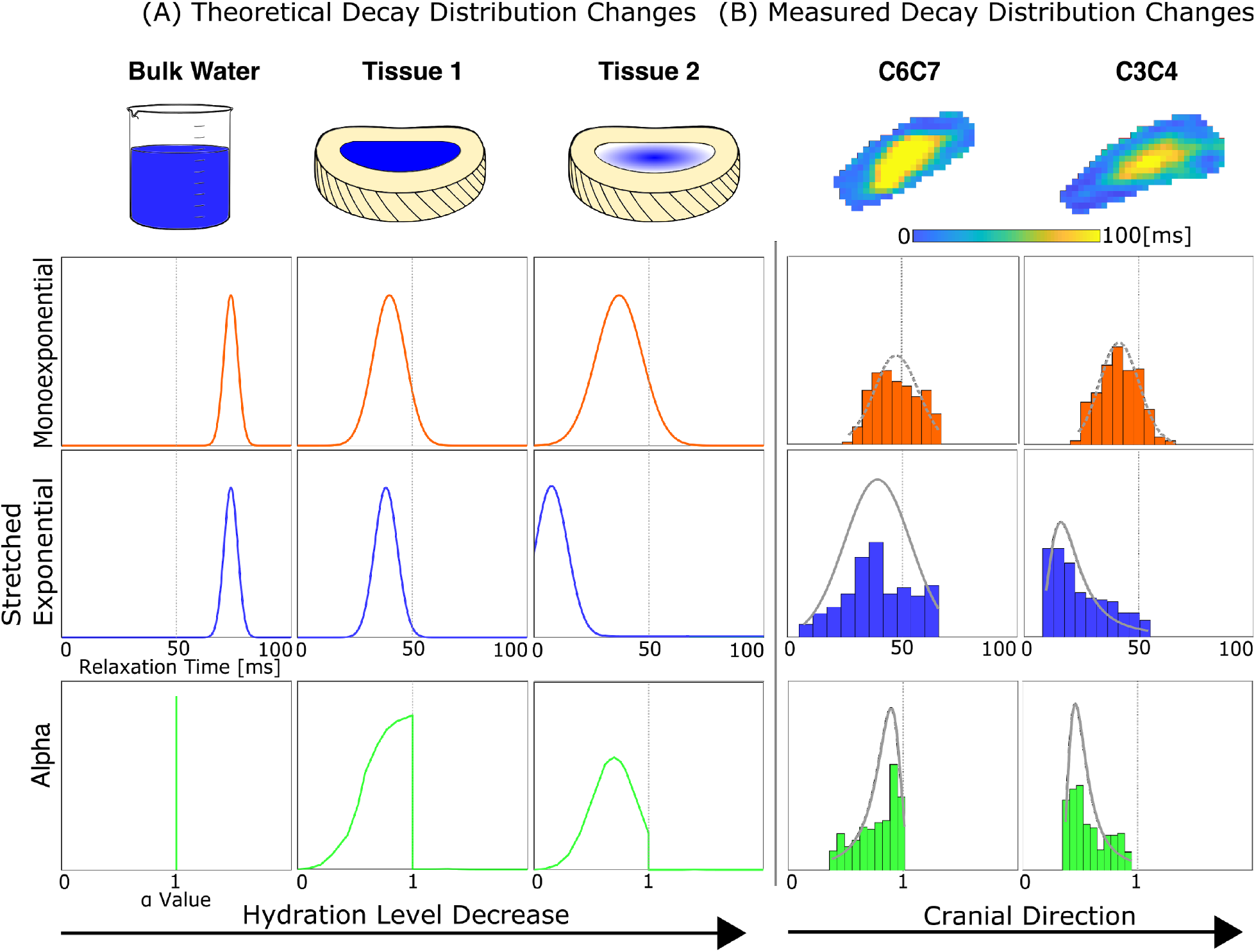
Anomalous relaxation behavior manifests due to increased tissue complexity and variation in the local microenvironment. (A) The monoexponential and SE models do not differ for the bulk water case (with α values all being one). Increasing sample complexity (e.g., due to changes in water content, pH, chemical exchange) leads to changes in relaxation values and dispersion, and the potential for higher sensitivity to compositional differences. (B) IVD levels, with variation in structure that depends on location in the spinal column, shows distinct differences in patterns of anomalous relaxation values, establishing a foundation to probe underlying mechanisms and time-course changes.

The SE model has a significantly lower MSE (*p*<0.001) which when coupled with a qualitative residual analysis (Supplemental Figure 2) suggests a non-monoexponential model may be a more appropriate decay fit. Furthermore, the SE model can recover a monoexponential decay fit with an α parameter of 1 (Figure 1A). The resulting SE qMRI maps show greater spatial heterogeneity compared to the monoexponential qMRI map (Figure 1B). This texture increase could reflect an improved sensitivity of these parameters to spatially varying composition and thus serve as useful biomarkers.

Inspection of the stable distributions from a single subject highlights differences in model fits (Figure 2). Aggregate T_1*ρ*Mono_ and T_2Mono_ over the entire disc show no discernable trend by level. The T_1*ρ*SE_ and T_2SE_ values show an increase in the lower IVDs while the aτi*p* and aτ2 values show a level-wise association throughout, particularly for ρ_T1*ρ*_. Single subject component analysis (NP vs. AF) reflects greater level-wise shifts in the NP SE parameter distributions supporting the proposed application of this model (Supplemental Figure 3). The level-wise increase in SE parameters – most visible in the T_1*ρ*_ analysis – may be indicative of the level-wise increase in GAG content of the NP (Supplemental Table 1, Supplemental Figure 4). Moreover, the distinct tissue microenvironment of the NP and AF, especially high GAG and collagen content, respectively, may differentially influence the level-wise analysis observed (32) (Supplemental Figure 4).

The population-level T_1*ρ*_ results are consistent with the hypothesis that δ values reflect the expected level-wise variation in whole disc GAG content (Figure 3). The conventional monoexponential T_1*ρ*_ decay model has been shown to correlate with GAG content (16). Additionally, T_1*ρ*_ within the NP values decrease with disc degeneration (24). However, the conventional fit may provide a limited representation of the relaxation features available as they relate to important matrix-water interactions limiting standard relaxometry measures sensitivity to characterize IVD tissue status. The T_1*ρ*SE_ and α_T1*ρ*_ show greater variation in the caudal direction, with the α_T1*ρ*_ being the more prominent of the two. An analysis of NP and AF individually (Figure 3B, Supplemental Figure 4), separates the seemingly bimodal α_T1*ρ*_ distribution and details a relationship between NP parameter values and disc level indicating the NP is a major factor in the observed level-wise variation based on the entire disc, supporting a potential correlation between α values and NP microstructure (Supplemental Table 1). The normalized population level-wise analysis indicates a strong monotonic relationship with level for α_T1*ρ*_ (*p*=0.003, ρ=0.404; Figure 3C) suggesting the SE model parameter may be more sensitive to compositional differences than conventional methods.

The T_2_ parameters show no significant correlation to IVD level (Figure 4), suggesting T_1*ρ*_ may be a more sensitive method to investigate IVD compositional differences. The qualitative distribution changes that do exist are led by the NP differences. Although not significant, the α_T2_ values show higher inter-IVD differences than T_2Mono_. The difference in T_1*ρ*_ and T_2_ is likely due to the increased sensitivity of T_1*ρ*_ to probe the slower molecular interactions of proteoglycans than T_2_ (14,15).

A stretched-exponential model of T_1*ρ*_ IVD relaxation could yield more sensitive insights into IVD health. Current literature suggests a strong correlation between absolute GAG content, IVD health, and monoexponential T_1*ρ*_ IVD relaxation. However, *in vivo* approaches are typically at the population level (e.g. with cohorts greater than 15 participants (33–36) and/or not IVD location specific (17), yielding limited utility at the individual patient level. A patient level correlation between relative IVD location and T_1*ρ*_ values has to the best of our knowledge never been demonstrated. With the current SE model, we demonstrate level-wise variation indicating a before unknown measure of sensitivity which could prove useful in detecting finer IVD compositional changes, such as those present in early IVDD.

In our patient data near the location of fusion, structural or degenerative changes in discs close to implant hardware are detected using the SE model that are not observed using monoexponential model (Figure 5). The monoexponential level-wise analysis indicates similar IVD composition for C2C3-C4C5 to that observed in healthy volunteers at those levels. Closer to the implant site, the SE model parameters deviate from the healthy data (Figure 5C-D), suggesting a change in IVD tissue integrity in C4C5. Although IVDs adjacent to the implant site are at an increased risk of accelerated degenerative changes after spinal fusion surgery (37,38), these T_1*ρ*_ weighted images were acquired five months post-fusion so are unlikely to represent recently developed degeneration due to the fusion procedure. However, it is not uncommon for patients to have multi-level degenerative changes outside of the fused segments that are pre-existing to the surgery. Thus, the observed deviation in SE parameters, particularly ρ_T1*ρ*_, might indicate a higher sensitivity to structural integrity in a patient with known degenerative changes. Further work is needed to evaluate this approach in the setting of adjacent disc disease.

Multiexponential (anomalous) modeling of T_2_ relaxation has previously been compared with quantitative biochemical composition of the IVD showing correlations in explant tissues with varying degree of degeneration (3). We observed no level-wise variation in SE T_2_ model parameters in our healthy population. This could be due to a lack of sensitivity of spin-spin relaxation to any level-wise variation in the IVD tissue properties. It could also be a result of the more limited acquisition parameters used for *in vivo* imaging (e.g. echo times, signal quality, etc). However, even if multiexponential T_2_ analysis were to provide a better representation of the underlying tissue microstructure, numerically it is less feasible to use in vivo due to the substantially greater demands on both the amount and quality of decay data (22) compared with the SE model. Future studies will compare structural properties in model tissue systems with these relaxation model parameters and investigate their relationship to IVD function (39,40). More detailed studies of IVD pathogenesis and damage are needed to investigate the efficacy of the SE fit for non-invasive characterization of IVDD *in vivo*.

## CONCLUSION

Using a stretched-exponential model may lead to improved sensitivity and possibly provide earlier detection of IVDD (Figure 7). The α parameter is thought to reflect a distribution of relaxation times and thus reflect interactions of water protons with the tissue microenvironment (21). An increase in hydrostatic pressure with increasing caudal disc level is due to increased disc load (32). The α_T1*ρ*_ values indicate a monotonic level-wise relationship consistent with reported variation in GAG concentration in healthy IVDs. Both SE model parameters in the patient data near the fusion site deviate from those observed at comparable levels in healthy subjects. The T_1*ρ*_ SE model may be a more sensitive analytical method for IVD compositional difference detection than the conventional model, holding potential as a biomarker for earlier IVDD detection.

**Figure 7:**
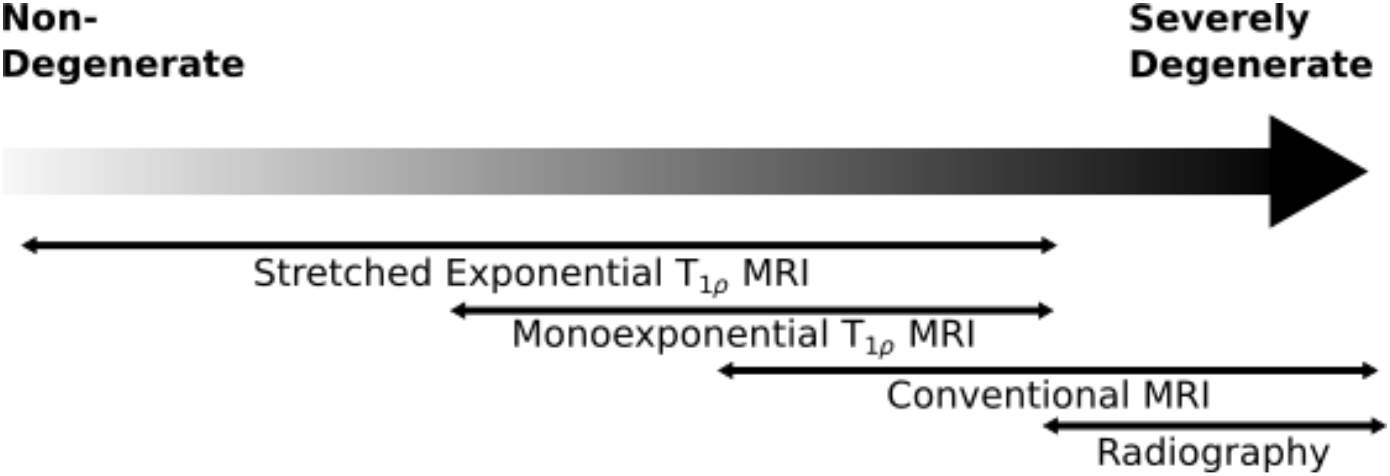
The larger dynamic range of the SE model compared to conventional methods may lead to earlier IVDD detection. The SE model’s increased dynamic range allows higher sensitivity to compositional differences, potentially detecting subtler IVD changes compared to conventional methods. The SE model could lead to improved biomarkers for screening and early detection of IVDD (Adapted from (17)).

## AUTHOR CONTRIBUTIONS

R.L.W., L.B., W.K., and D.A.R performed the experiments described. R.L.W, C.P.N., and D.A.R. conceived of the study, and designed the experiments. R.L.W. wrote the manuscript. All authors edited and reviewed the manuscript.

## ACKNOWLEDGEMENTS

We appreciate funding support from NIH grants R01 AR063712 and R21 AR066665. Additionally, research reported in this publication was supported by the National Institute of Biomedical Imaging and Bioengineering of the National Institutes of Health under award Number P41-EB015893.

## SUPPLEMENTAL TABLES

**Supplemental Table 1:** The T_1*ρ*_ values for each fitting parameter tissue compartment by level with significance and Spearman’s ρ. The level-wise analysis indicates significant trends in all compartments of the α_T1*ρ*_ data and the T_1*ρ*SE_ nucleus pulposus data indicating the SE model may be a more sensitive analysis method of IVD composition. Data is presented as mean ± standard error of the mean. * indicates *p*<0.05.

| Model Parameter | Tissue Compartment | IVD $\delta$ Value | | | | | $p$ value | Spearman's $\rho$ |
| --- | --- | --- | --- | --- | --- | --- | --- | --- |
|  |  | C2C3 | C3C4 | C4C5 | C5C6 | C6C7 |  |  |
| $T_{1\rho Mono}$ [ms] | Whole Disc | 46.19<br>$\pm 1.85$ | 44.69±<br>2.29 | 47.62<br>$\pm 1.62$ | 42.17<br>$\pm 1.56$ | 42.06<br>$\pm 1.32$ | 0.20 | -0.18 |
| | Nucleus Pulposus | 46.75<br>$\pm 4.49$ | 51.46±<br>2.81 | 52.95<br>$\pm 2.59$ | 48.77<br>$\pm 2.14$ | 54.98<br>$\pm 2.01$ | 0.79 | 0.035 |
| | Annulus Fibrosus | 39.99<br>$\pm 3.50$ | 43.37<br>$\pm 2.57$ | 44.20<br>$\pm 1.39$ | 39.30<br>$\pm 1.53$ | 40.52<br>$\pm 1.36$ | 0.31 | -0.14 |
| $T_{1\rho SE}$ [ms] | Whole Disc | 25.61<br>$\pm 3.03$ | 25.29±<br>3.15 | 33.44<br>$\pm 2.57$ | 28.69<br>$\pm 2.88$ | 27.92<br>$\pm 2.76$ | 0.32 | 0.14 |
| | Nucleus Pulposus | 34.80<br>$\pm 4.28$ | 36.42<br>$\pm 3.67$ | 43.47<br>$\pm 3.96$ | 40.39<br>$\pm 3.64$ | 48.74<br>$\pm 3.29$ | 0.021* | 0.31 |
| | Annulus Fibrosus | 18.90<br>$\pm 2.67$ | 20.96<br>$\pm 2.46$ | 26.64<br>$\pm 2.03$ | 24.22<br>$\pm 2.36$ | 23.90<br>$\pm 2.73$ | 0.23 | 0.17 |
| $\alpha_{T1\rho}$ | Whole Disc | 0.52<br>$\pm 0.034$ | 0.53<br>$\pm 0.035$ | 0.57<br>$\pm 0.035$ | 0.63<br>$\pm 0.049$ | 0.68<br>$\pm 0.048$ | 0.0025* | 0.40 |
| | Nucleus Pulposus | 0.57<br>$\pm 0.047$ | 0.60<br>$\pm 0.044$ | 0.66<br>$\pm 0.046$ | 0.68<br>$\pm 0.057$ | 0.80<br>$\pm 0.056$ | 0.0029* | 0.39 |
| | Annulus Fibrosus | 0.47<br>$\pm 0.022$ | 0.52<br>$\pm 0.033$ | 0.53<br>$\pm 0.032$ | 0.56<br>$\pm 0.049$ | 0.60<br>$\pm 0.043$ | 0.017* | 0.32 |

## SUPPLEMENTAL FIGURES

**Supplemental Figure 1:**
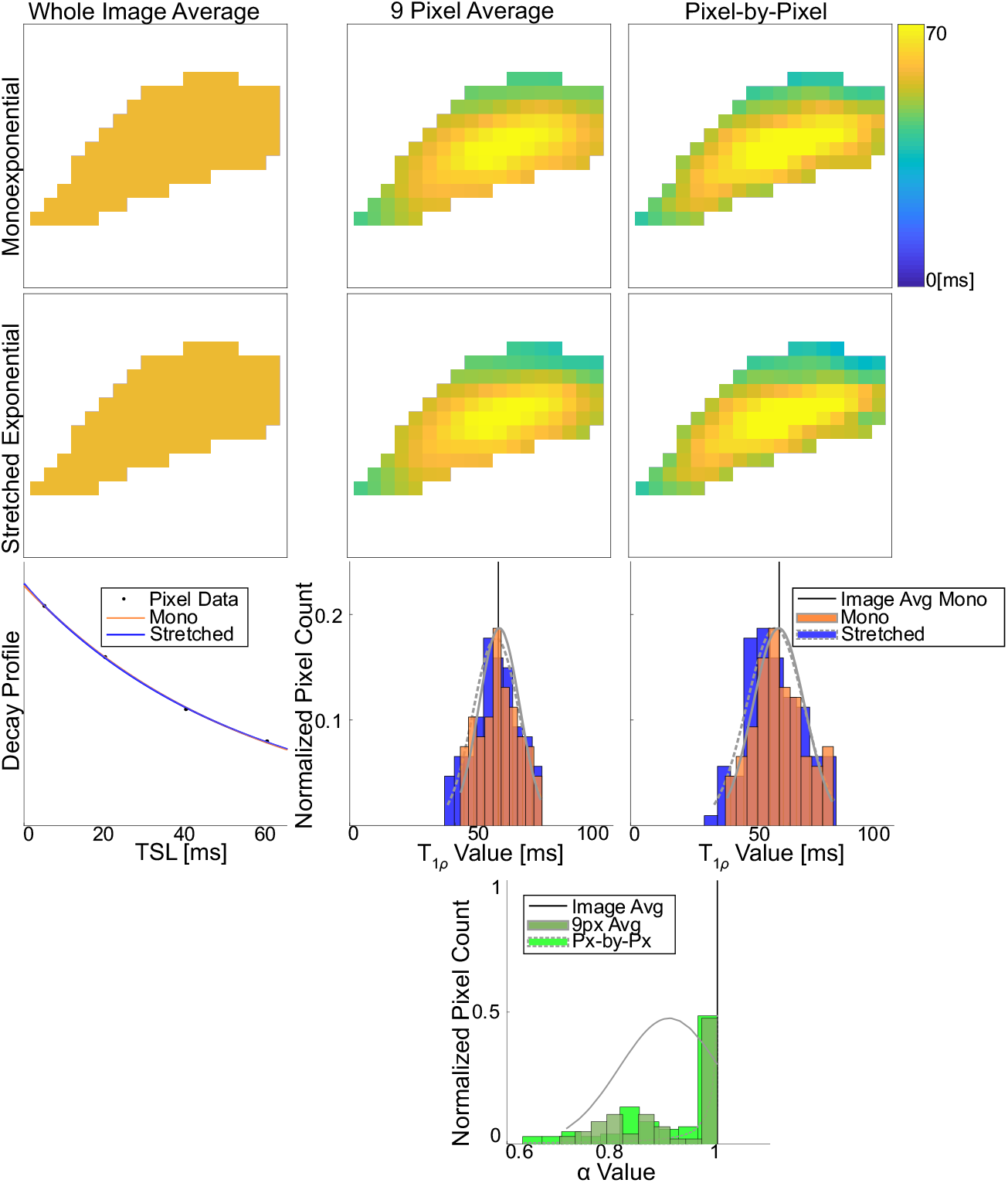
A pixel-by-pixel processing method is a reliable processing method, allowing for maximum spatial information retention. Evaluating the entire disc (by averaging all values at each time point) is not a spatially sensitive diagnostic tool, yet it provides an excellent baseline for measuring other processing methods due to its inherent noise minimization. Whole image T_1*ρ*_ averages for monoexponential and SE models are 56.4 ms and 56.2 ms, respectively. The 9-pixel moving window average allows for increased spatial information with T_1*ρ*_ δ values of 56.3 ms and 54.9 ms for monoexponential and SE respectively. However, the pixel-by-pixel method was chosen for analysis as its T_1*ρ*_ δ values (Monoexponential: 56.8 ms SE: 55.6 ms) were closer to that of the whole image average than the 9-pixel method suggesting minimal noise while retaining a higher level of spatial detail.

**Supplemental Figure 2:**
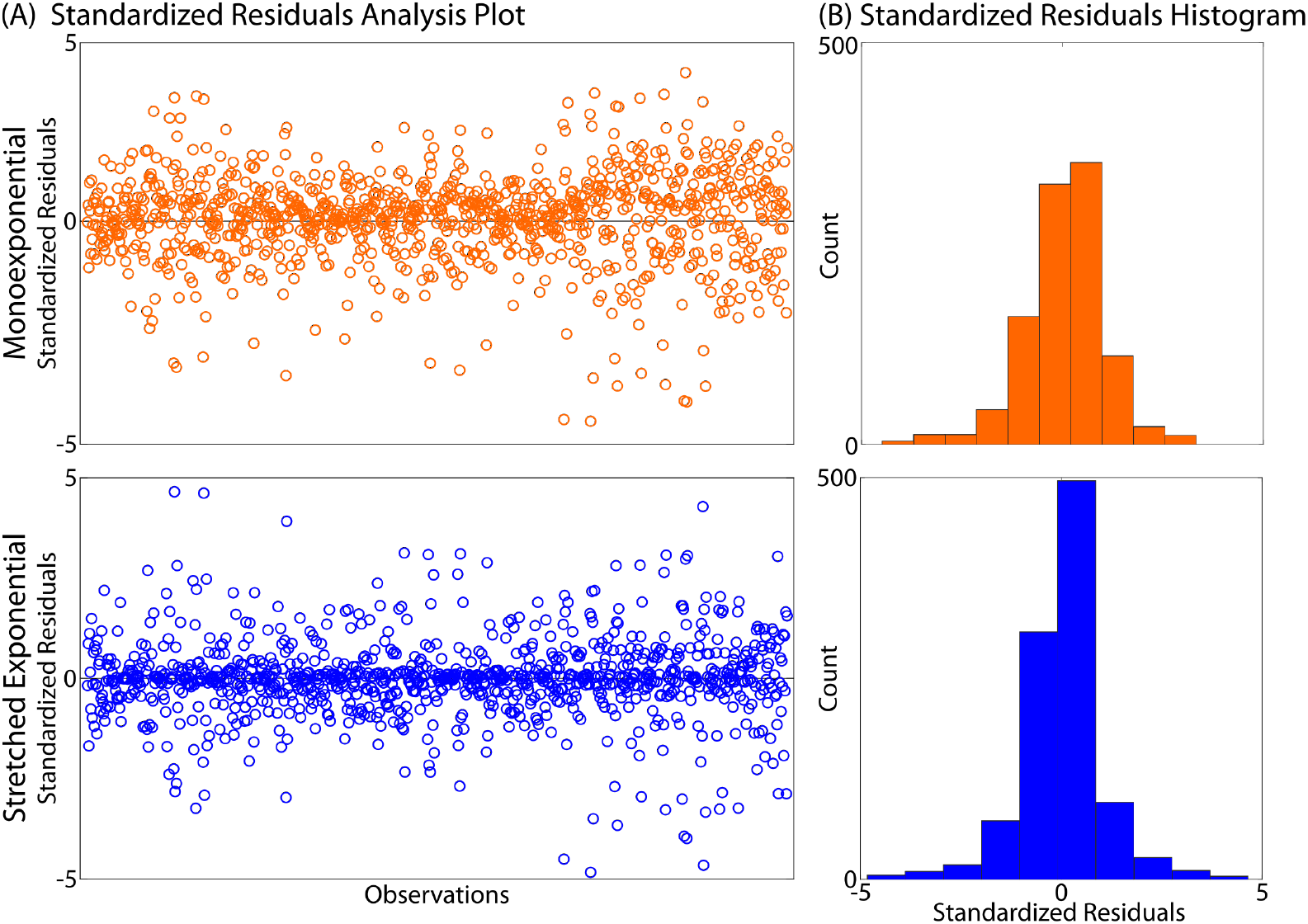
Standardized residual analysis indicates similar levels of fit for the monoexponential and stretched-exponential decay functions. (A) Standardized residuals from monoexponential and stretched-exponential fits of each TSL from all pixels in a single IVD. (B) Histograms of the monoexponential and stretched-exponential fit standardized residuals. The residual plots detail randomly distributed standardized residuals while the histograms indicate a normal distribution with minimal skewness.

**Supplemental Figure 3:**
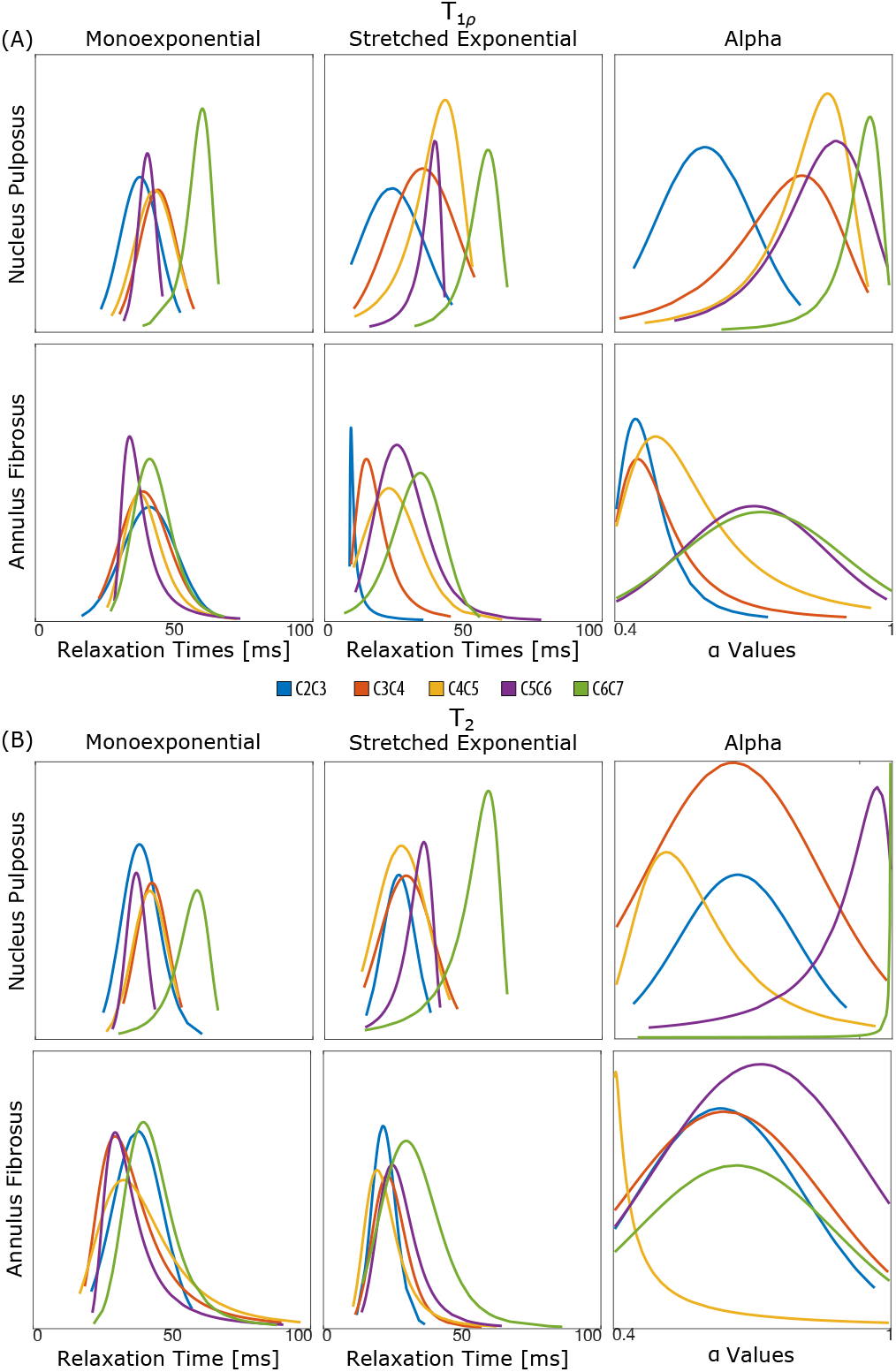
Single-subject nucleus pulposus T_1*ρ*_ and α_T1*ρ*_ stable distributions show a qualitative correspondence with IVD level, whereas all others do not. (A) The SE T_1*ρ*_ stable distributions show an apparent level-wise trend in the NP while monoexponential NP and all AF distributions do not (B) The T_2_ stable distributions show no trend.

**Supplemental Figure 4:**
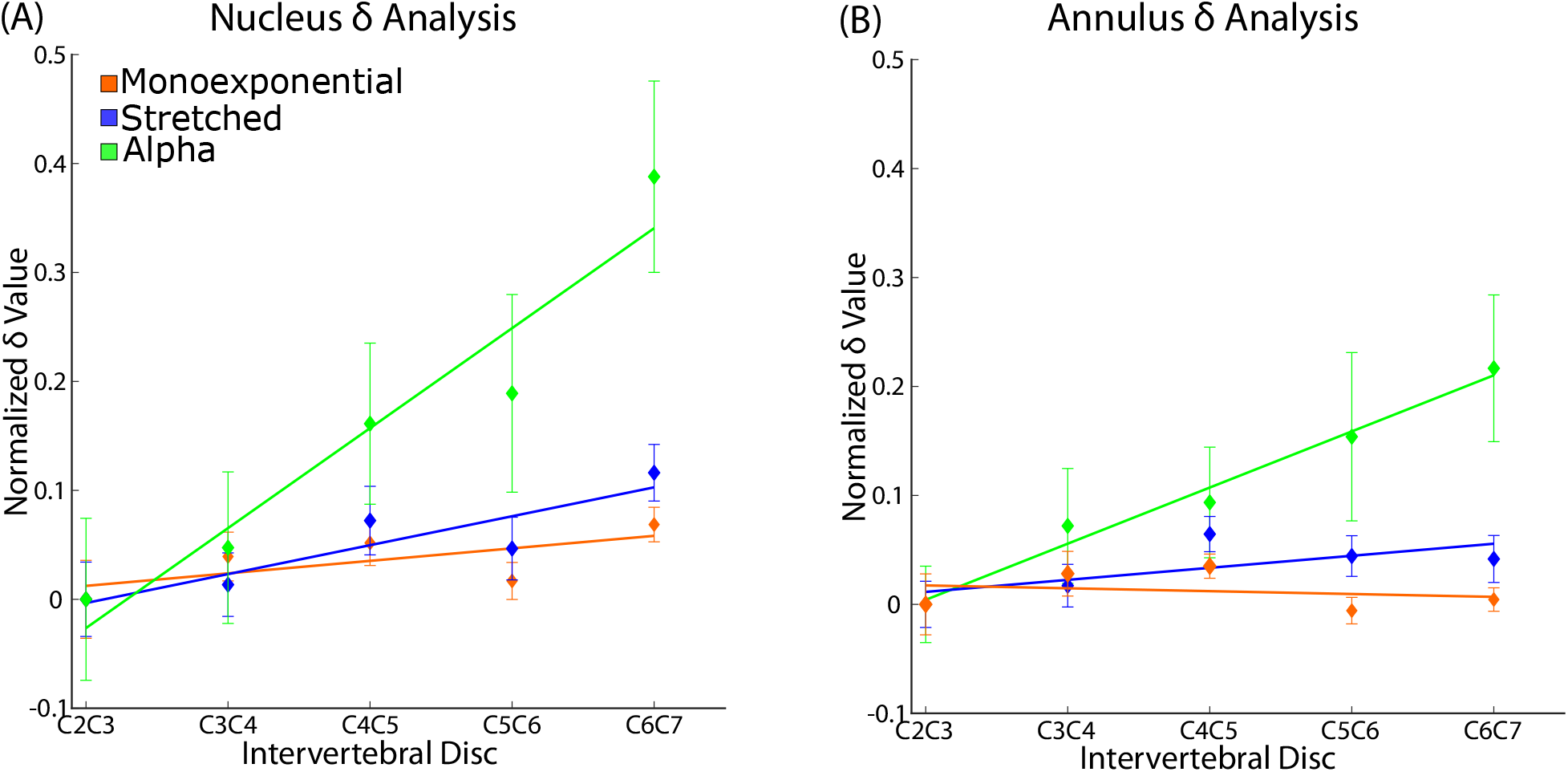
T_1*ρ*_ compartmental data suggests a more sensitive monotonic relationship in the nucleus pulposus than the annulus fibrosus. (A) The nucleus pulposus level-wise analysis presents a significant relationship with position for T_1*ρ*SE_ and a with no monotonic relationship in the T_1*ρ*Mono_ data (Supplemental Table 1). (B) The annulus fibrosus level-wise analysis details a significant relationship in the α data only. The stronger monotonic relationship in the level-wise nucleus pulposus analysis, as evidenced by the difference in *ρ* values, indicates that whole disc level-wise changes are driven by nucleus pulposus composition.

## REFERENCES

1. Adams MA, Roughley PJ. What is Intervertebral Disc Degeneration, and What Causes It? Spine (Phila. Pa. 1976). 2006;31:2151–2161 doi: 10.1097/01.brs.0000231761.73859.2c.

2. Martin BI, Deyo RA, Mirza SK, et al. Expenditures and Health Status Among Adults With Back and Neck Problems. JAMA 2008;299:656 doi: 10.1001/jama.299.6.656.

3. Nightingale T, MacKay A, Pearce RH, Whittall KP, Flak B. A model of unloaded human intervertebral disk based on NMR relaxation. Magn. Reson. Med. 2000;43:34–44 doi: 10.1002/(SICI)1522-2594(200001)43:1<34::AID-MRM5>3.0.CO;2-7.

4. Chen C, Huang M, Han Z, et al. Quantitative T2 Magnetic Resonance Imaging Compared to Morphological Grading of the Early Cervical Intervertebral Disc Degeneration: An Evaluation Approach in Asymptomatic Young Adults. PLoS One 2014;9 doi: 10.1371/journal.pone.0087856.

5. Shapiro F, Koide S, Glimcher MJ. Cell origin and differentiation in the repair of full-thickness defects of articular cartilage. J. Bone Joint Surg. Am. 1993;75:532–53.

6. Gilbert SJ, Singhrao SK, Khan IM, et al. Enhanced Tissue Integration During Cartilage Repair *In Vitro* Can Be Achieved by Inhibiting Chondrocyte Death at the Wound Edge. Tissue Eng. Part A 2009; 15:1739–1749 doi: 10.1089/ten.tea.2008.0361.

7. Xue J, He A, Zhu Y, et al. Repair of articular cartilage defects with acellular cartilage sheets in a swine model. Biomed. Mater. 2018;13:025016 doi: 10.1088/1748-605X/aa99a4.

8. Chan DD, Khan SN, Ye X, et al. Mechanical Deformation and Glycosaminoglycan Content Changes in a Rabbit Annular Puncture Disc Degeneration Model. Spine (Phila. Pa. 1976). 2011;36:1438–1445 doi: 10.1097/BRS.0b013e3181f8be52.

9. Crisco JJ, McGovern RD, Wolfe SW. Noninvasive technique for measuringin vivo three-dimensional carpal bone kinematics. J. Orthop. Res. 1999;17:96–100 doi: 10.1002/jor.1100170115.

10. Hodge WA, Fijan RS, Carlson KL, Burgess RG, Harris WH, Mann RW. Contact pressures in the human hip joint measured in vivo. Proc. Natl. Acad. Sci. U. S. A. 1986;83:2879–83.

11. Nieminen MT, Töyräs J, Rieppo J, et al. Quantitative MR microscopy of enzymatically degraded articular cartilage. Magn. Reson. Med. 2000;43:676–681 doi: 10.1002/(SICI)1522-2594(200005)43:5<676::AID-MRM9>3.0.CO;2-X.

12. Lüsse S, Claassen H, Gehrke T, et al. Evaluation of water content by spatially resolved transverse relaxation times of human articular cartilage.

13. Liess C, Lü Sse S, Karger N, Heller M, Glü C-C. Detection of changes in cartilage water content using MRI T 2-mapping in vivo. Osteoarthr. Cartil. 2002;10:907–913 doi: 10.1053.

14. Akella S V, Regatte RR, Gougoutas AJ, et al. Proteoglycan-induced changes in T1rho-relaxation of articular cartilage at 4T. Magn. Reson. Med. 2001;46:419–23.

15. Regatte RR, Akella SVS, Borthakur A, Reddy R. Proton spin-lock ratio imaging for quantitation of glycosaminoglycans in articular cartilage. J. Magn. Reson. Imaging 2003;17:114–21 doi: 10.1002/jmri.10228.

16. Johannessen W, Auerbach JD, Wheaton AJ, et al. Assessment of human disc degeneration and proteoglycan content using T1rho-weighted magnetic resonance imaging. Spine (Phila. Pa. 1976). 2006;31:1253–7 doi: 10.1097/01.brs.0000217708.54880.51.

17. Auerbach JD, Johannessen W, Borthakur A, et al. In vivo quantification of human lumbar disc degeneration using T1p-weighted magnetic resonance imaging. Eur. Spine J. 2006;15:338–344 doi: 10.1007/s00586-006-0083-2.

18. Duvvuri U, Reddy R, Patel SD, Kaufman JH, Kneeland JB, Leigh JS. T1p-relaxation in articular cartilage: Effects of enzymatic degradation. Magn. Reson. Med. 1997;38:863–867 doi: 10.1002/mrm.1910380602.

19. Regatte RR, Akella SVS, Borthakur A, Kneeland JB, Reddy R. Proteoglycan depletion-induced changes in transverse relaxation maps of cartilage: comparison of T2 and T1rho. Acad. Radiol. 2002;9:1388–94.

20. Menezes NM, Gray ML, Hartke JR, Burstein D. T2 andT1rho MRI in articular cartilage systems. Magn. Reson. Med. 2004;51:503–509 doi: 10.1002/mrm.10710.

21. Reiter DA, Magin RL, Li W, Trujillo JJ, Pilar Velasco M, Spencer RG. Anomalous T2 relaxation in normal and degraded cartilage. Magn. Reson. Med. 2016;76:953–962 doi: 10.1002/mrm.25913.

22. Reiter DA, Roque RA, Lin P-C, et al. Mapping proteoglycan-bound water in cartilage: Improved specificity of matrix assessment using multiexponential transverse relaxation analysis. Magn. Reson. Med. 2011;65:377–384 doi: 10.1002/mrm.22673.

23. June RK, Neu CP, Barone JR, Fyhrie DP. Polymer mechanics as a model for short-term and flow-independent cartilage viscoelasticity. Mater. Sci. Eng. C 2011;31:781–788 doi: 10.1016/J.MSEC.2010.11.029.

24. Paul CPL, Smit TH, de Graaf M, et al. Quantitative MRI in early intervertebral disc degeneration: T1rho correlates better than T2 and ADC with biomechanics, histology and matrix content Nikitovic D, editor. PLoS One 2018;13:e0191442 doi: 10.1371/journal.pone.0191442.

25. Magin RL, Li W, Pilar Velasco M, et al. Anomalous NMR relaxation in cartilage matrix components and native cartilage: fractional-order models. J. Magn. Reson. 2011;210:184–91 doi: 10.1016/j.jmr.2011.03.006.

26. Santyr GE, Henkelman RM, Bronskill MJ. Variation in measured transverse relaxation in tissue resulting from spin locking with the CPMG sequence. J. Magn. Reson. 1988;79:28–44 doi: 10.1016/0022-2364(88)90320-4.

27. Li X, Wyatt C, Rivoire J, et al. Simultaneous acquisition of T_1p_ and T_2_ quantification in knee cartilage: Repeatability and diurnal variation. J. Magn. Reson. Imaging 2014;39:1287–1293 doi: 10.1002/jmri.24253.

28. Wirth W, Maschek S, Roemer FW, Eckstein F. Layer-specific femorotibial cartilage T2 relaxation time in knees with and without early knee osteoarthritis: Data from the Osteoarthritis Initiative (OAI). Nat. Publ. Gr. 2016 doi: 10.1038/srep34202.

29. Chan DD, Cai L, Butz KD, et al. Functional MRI can detect changes in intratissue strains in a full thickness and critical sized ovine cartilage defect model. J. Biomech. 2018;66:18–25 doi: 10.1016/J.JBIOMECH.2017.10.031.

30. Witschey WRT, Borthakur A, Elliott MA, et al. Artifacts in T1ρ-weighted imaging: Compensation for B1 and B0 field imperfections. J. Magn. Reson. 2007;186:75–85 doi: 10.1016/j.jmr.2007.01.015.

31. Bloch F. Nuclear Induction. Phys. Rev. 1946;70:460–474 doi: 10.1103/PhysRev.70.460.

32. Schleich C, Müller-Lutz A, Zimmermann L, et al. Biochemical imaging of cervical intervertebral discs with glycosaminoglycan chemical exchange saturation transfer magnetic resonance imaging: feasibility and initial results. Skeletal Radiol. 2016;45:79–85 doi: 10.1007/s00256-015-2251-0.

33. Blumenkrantz G, Zuo J, Li X, Kornak J, Link TM, Majumdar S. In vivo 3.0-tesla magnetic resonance T1ρ and T2 relaxation mapping in subjects with intervertebral disc degeneration and clinical symptoms. Magn. Reson. Med. 2010;63:1193–1200 doi: 10.1002/mrm.22362.

34. Pandit P, Talbott JF, Pedoia V, Dillon W, Majumdar S. T1ρ and T2-based characterization of regional variations in intervertebral discs to detect early degenerative changes. J. Orthop. Res. 2016;34:1373–1381 doi: 10.1002/jor.23311.

35. Borthakur A, Maurer PM, Fenty M, et al. T1ρ magnetic resonance imaging and discography pressure as novel biomarkers for disc degeneration and low back pain. Spine (Phila. Pa. 1976). 2011;36:2190–6 doi: 10.1097/BRS.0b013e31820287bf.

36. Zuo J, Joseph GB, Li X, et al. In vivo intervertebral disc characterization using magnetic resonance spectroscopy and T1ρ imaging: association with discography and Oswestry Disability Index and Short Form-36 Health Survey. Spine (Phila. Pa. 1976). 2012;37:214–21 doi: 10.1097/BRS.0b013e3182294a63.

37. Akamaru T, Kawahara N, Tim Yoon S, et al. Adjacent segment motion after a simulated lumbar fusion in different sagittal alignments: a biomechanical analysis. Spine (Phila. Pa. 1976). 2003;28:1560–6.

38. Hilibrand AS, Robbins M. Adjacent segment degeneration and adjacent segment disease: the consequences of spinal fusion? Spine J. 2004;4:S190–S194 doi: 10.1016/J.SPINEE.2004.07.007.

39. Chan DD, Neu CP. Intervertebral disc internal deformation measured by displacements under applied loading with MRI at 3T. Magn. Reson. Med. 2014;71:1231–1237 doi: 10.1002/mrm.24757.

40. Chan DD, Gossett PC, Butz KD, Nauman EA, Neu CP. Comparison of intervertebral disc displacements measured under applied loading with MRI at 3.0 T and 9.4 T. J. Biomech. 2014;47:2801–2806 doi: 10.1016/J.JBIOMECH.2014.05.026.

